# Selection of an Ideal Machine Learning Framework for Predicting Perturbation Effects on Network Topology of Bacterial KEGG Pathways

**DOI:** 10.1101/2022.07.21.501034

**Authors:** Michael Robben, Mohammad Sadegh Nasr, Avishek Das, Manfred Huber, Justyn Jaworski, Jon Weidanz, Jacob Luber

## Abstract

Biological networks for bacterial species are used to assign functional information to newly sequenced organisms but network quality can be largely affected by poor gene annotations. Current methods of gene annotation use homologous alignment to determine orthology, and have been shown to degrade network accuracy in non-model bacterial species. To address these issues in the KEGG pathway database, we investigated the ability for machine learning (ML) algorithms to re-annotate bacterial genes based on motif or homology information. The majority of the ensemble, clustering, and deep learning algorithms that we explored showed higher prediction accuracy than CD-hit in predicting EC ID, Map ID, and partial Map ID. Motif-based, machine-learning methods of annotation in new species were more accurate, faster, and had higher precision-recall than methods of homologous alignment or orthologous gene clustering. Gradient boosted ensemble methods and neural networks also predicted higher connectivity of networks, finding twice as many new pathway interactions than blast alignment. The use of motif-based, machine-learning algorithms in annotation software will allow researchers to develop powerful network tools to interact with bacterial microbiomes in ways previously unachievable through homologous sequence alignment.

**CCS CONCEPTS:** **• Applied computing** → **Computational biology**; **Life and medical sciences**; **Bioinformatics**; • **Computing methodologies** → **Machine learning algorithms**; **Machine learning approaches**.

**ACM Reference Format:** Michael Robben, Mohammad Sadegh Nasr, Avishek Das, Manfred Huber, Justyn Jaworski, Jon Weidanz, and Jacob Luber. 2022. Selection of an Ideal Machine Learning Framework for Predicting Perturbation Effects on Network Topology of Bacterial KEGG Pathways. In *The 13th ACM Conference on Bioinformatics, Computational Biology, and Health Informatics, August 07–10, 2022, Chicago, IL*. ACM, New York, NY, USA, 11 pages. https://doi.org/XXXXXXX.XXXXXXX

## 1 INTRODUCTION

The human gut microbiome is inextricably linked to human health, contributing to the progression of inflammatory disease [25], neurological conditions [20], and even cancer [18, 62]. Since the advent of large scale genome sequencing, many projects have sought to sequence and characterize the bacteria found colonizing the human gut system [19]. Much of the work done to characterize these species has focused on taxonomic diversity analysis through 16S sequencing, but more studies now perform genome assembly or metatranscriptomics to ascribe functional relevance to populations and diseases [11, 14, 32]. Such studies have been known to assemble 100’s to 1000’s of bacterial genomes per sequencing run, which requires a significant amount of computational power and high quality functional information to annotate [1].

Many large database networks hold annotation information for microbial genes such as PFAM [34], Interpro [4], Uniprot [54], SEED [38], and the Refseq database [37]. The KEGG database contains curated information on protein pathways vital to the cellular function of gut bacteria [22]. Enrichment of specific KEGG pathways have been previously associated with disease prevalence [26, 27, 56]. However, pathway annotations in the database are notoriously poor, with surveys of sequenced species showing proteomic annotation as low as 14% in some cases [16, 33, 47]. This disparity is largest in evolved taxa furthest from well characterized species such as *Escherichia coli* and *Bacillus subtilis*. This issue of quality in biological networks is currently of major concern in the field of network biology. Many well curated networks remain incomplete with as much as 80% missing data in protein-protein interaction networks [17]. The effects of network completeness from poor annotation can be seen between a model bacterial organism like *E. coli* and a non-model gut microbe like *Streptococcus macedonicus* (Fig 1a). Recent reports show how poorly curated data can effect network performance, such as how a lack of curation of the blast nr database has resulted in misclassification of disease causing microbes [3]. It stands that in order to better understand the role of the gut microbiome in human disease, we first need to improve the quality of protein annotations.

**Figure 1:**
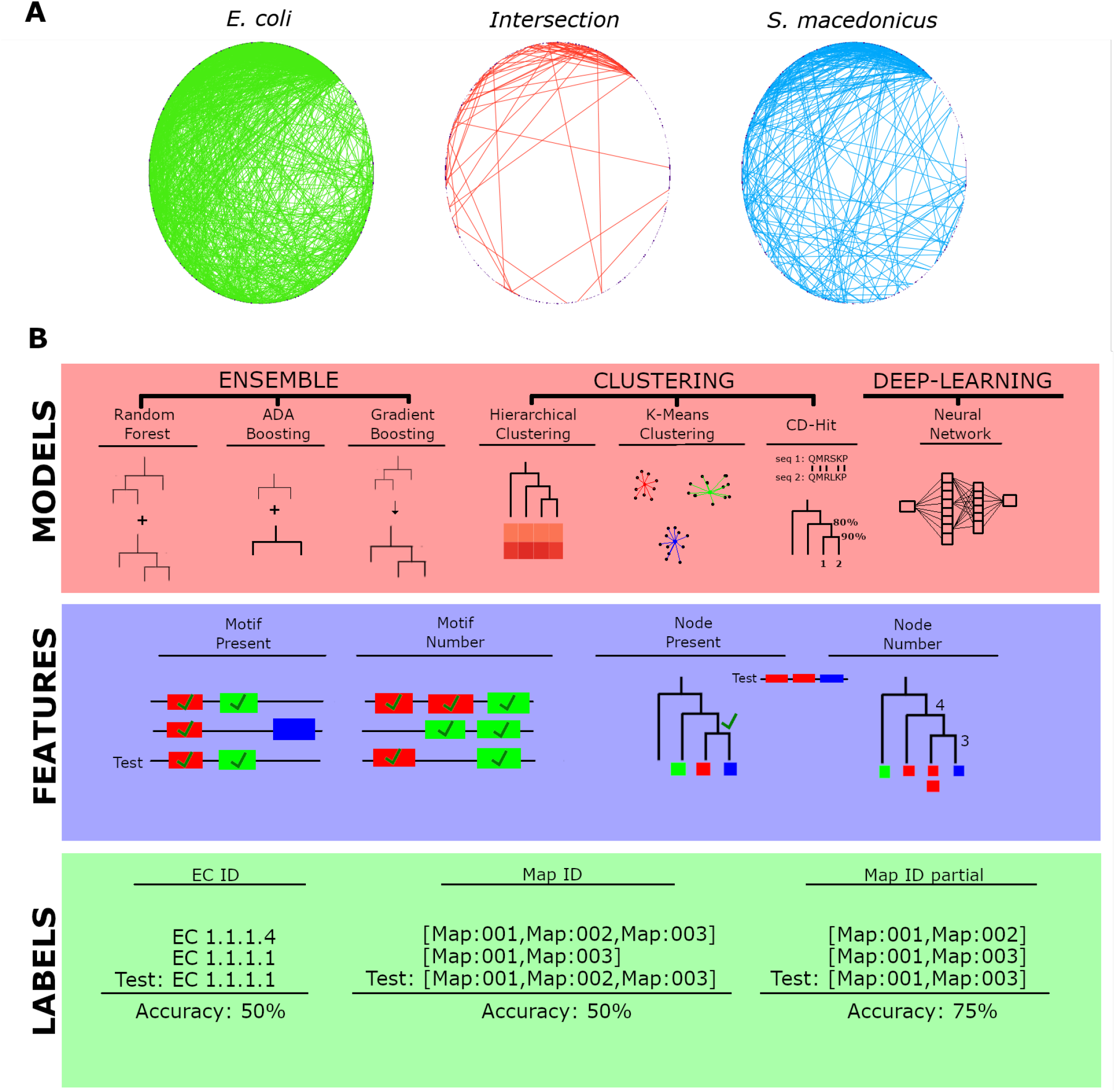
The effect of poor annotation quality on KEGG pathway networks and the strategy for benchmark analysis of machine learning algorithms. (A) Edge node circle diagram shows the effect of lack of annotation depth on network topology for KEGG pathways. Edges unique to *E. coli* are displayed in red, edges unique to *S. macedonicus* are displayed in blue, and edges common between both species are shown in yellow. (B) The three ensemble methods use multiple learners to make separate classifications on sampled data and then aggregate the most common classifier as the predicted set. In comparison, clustering is an unsupervised algorithm that groups similar data points to predict commonalities between them. We used aggregation of predictors within each cluster to classify the test data set in order to compare performance against supervised methods. A neural network was used to benchmark against the 6 classifiers to understand how classical machine learning algorithms could perform in comparison to a deep-learning algorithm. Four feature sets were trained on the models to determine the effect that motif based and homology based information has on the outcome of each learner. Each model was asked to predict 3 different labels given a test feature set, Enzyme Commission number (EC ID), a string of KEGG pathway identifiers (Map ID) and individual Map IDs specific to each gene (Partial Map ID).

Currently, the most widely used method to annotate newly sequenced bacterial genes is BLAST (Basic Local Alignment Search Tool) [45] alignment of proteins to previously assembled genomes [50]. New sequence alignment algorithms like Diamond [6], domain based alignment algorithms like HMMer [9], and nucleotide k-mer alignment algorithms like kraken2 [58] have been developed to improve the speed and fidelity of annotation. However, these traditional methods lead to the problem that newly sequenced genomes are annotated by alignment to other newly sequenced genomes and the loss of functional information compounds as more genomes are assembled [36]. To reduce this loss of data, some groups have published new methods of annotation, such as CD-hit [30], that work by clustering orthologous gene groups and can be applied to entire databases. Orthologous clusters of bacterial Open Reading Frames (ORFs) may retain functional similarity at rates greater than 95% and thus can be used to assign functional relation to newly predicted genes [28, 29]. Although homologous alignment of protein or DNA sequence is the norm for bacterial gene annotations, it has been shown that alignment to Hidden Markov Models (HMMs) describing conserved domains results in higher annotation completeness [33]. At the same time, new programs such as the meme-suite [2], have been published to identify functional subunits of proteins and DNA, called motifs, that are found ubiquitous to a cadre of proteins agnostic of predescribed function.

Machine learning algorithms are the ideal methods of computational classification, and have been successfully applied to the field of disease prediction [53]. A few studies have explored the ability for machine learning to functionally annotate bacterial genes, specifically through naive bayes classifiers [44], support vector machines [55], and deep learning neural networks [31]. A classification model that has not been investigated yet are ensemble methods, which have shown high success for classification in large, multi-class problems from imbalanced data sets [24]. We sought to compare ensemble, clustering, and deep learning models on their ability to predict Enzyme Commision (EC) identifiers and Map IDs from the KEGG database. We also pursued the efficacy of using protein motifs as features for the models in comparison to sequence homology in an effort to increase prediction accuracy and reduce computational time.

## 2 METHODS

### 2.1 Data Mining and Curation

Data from the KEGG database was queried by custom python script to obtain information on all genes from species of interest through the KEGG Rest API. The following identifiers were retrieved by “gene_id”: “ko_id”, “ec_id”, ”map_id”, “aa_seq”, and “nt_seq”, and were output as a comma delineated table. We queried the database for genes from *E. coli, Bacteroides intestinalis*, and *S. macedonicus* and subsetted only genes with annotations to KEGG pathways.

### 2.2 Feature Preparation

Protein sequences from pathway annotated genes were aligned by multiple sequence alignment through the MAFFT program [23]. Resulting alignments were then used to construct a phylogenetic tree using the neighbor joining algorithm with the phangorn package in R [46]. Motifs were predicted in all query genes through the meme program and resulting HMM models were used to identify motifs in each gene through fimo [15]. A custom R script was used to construct feature sets from motif hit files and phylogenetic tree data with the assistance of the tidytree R package [59]. We utilized 4 different feature sets named “Motif_present”, “Motif_number”, “Nodes_present”, and “Nodes_number”. Node based feature sets utilize phylogenetic alignments of genes from the training data but could be reconstructed solely from motif alignments, representing an increase in performance speed over traditional annotation techniques that rely on homologous sequence alignment.

### 2.3 Machine Learning Algorithms

Machine learning algorithms were compared for accuracy in their ability to predict functional annotation in bacterial genes using various feature sets. All classifiers were built to predict a single class from a multi-class problem using 2 different sets of labels, “ec_id” and “map_id”. In order to calculate prediction accuracy of labels we used the following formula:

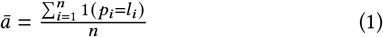

Where the accuracy is equal to the sum of predictions *p* that match the true string *l* as a boolean divided by the *n* number of test genes. Because Map IDs are actually a string of multiple Map IDs we also predicted partial accuracy of map ID prediction:

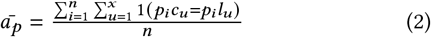

The accuracy of partial map IDs is equal to the average of the number of Map IDs *c* within each predicted string *p* that match an identifier *l* divided by the *n* number of test genes.

We assessed metrics that could be determined and compared between all models. True positive (3), false positive (4), true negative (5), and false negatives (6) were computed from confusion matrices using the following algorithms.

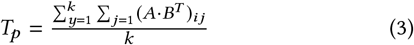

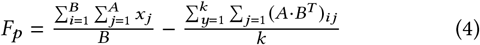

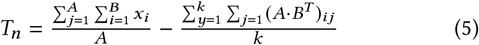

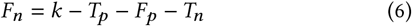

Where *k* is the total number of classes, *A* and *B* are the column and row dimensions, *i* and *j* are iterators of columns and rows respectively. Precision (7), recall (8), false positive rate (9), and log loss (10) were calculated for each machine learning algorithm.

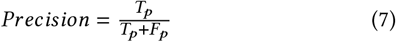

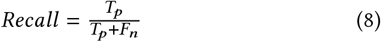

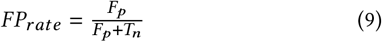

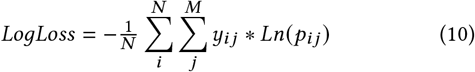

Where *N* is the number of genes, *M* is the number of different labels, *y*_*i j*_ is the binary variable with the expected labels and *p*_*i j*_ is the classification probability output by the classifier for the *i*-th instance and *j*-th label. Genes from *E. coli* and *B. intestinalis* were used as training data for the models. Models were trained on 75% of the training data and tested against 25% of the training data for validation. When tested against genes from *S. macedonicus* we trained the models on all of the training data.

#### 2.3.1 Random Forest Classifier

We implemented a random forest [5] classifier through the sklearn python library. Random Forest algorithms apply multiple decision trees to solve classification problems and then use ensemble voting to determine the most common predicted label from all trees. Hyper parameterization led us to use the following parameters in all models: *‘n_estimators’ = 5000, ‘min_samples_split’ = 2, ‘min_samples_leaf’ = 1, ‘max_features’ = sqrt’, ‘max_depth’ = 10’*. Hyper parameterization was conducted with 5 fold cross validation.

#### 2.3.2 AdaBoost Classifier

The ada boost classifier [12] was implemented through the sklearn python library. Ada boosting works similar to random forests, but uses a forest of weak learners, often called stumps, and weighs each weak learner by contribution to accurate prediction. We used the following parameters as a result of hyper parameterization: *‘learning_rate’ = 0.4, ‘n_estimators’ = 7500, ‘max_depth’ = 9*. Hyper parameterization was conducted with 5 fold cross validation.

#### 2.3.3 Histogram Gradient Boosting Classifier

Histogram gradient boosting algorithms provide the iterative weighting of a gradient boosting algorithm in a fast histogram based learner algorithm [13]. We implemented the Histogram gradient boosting through the sklearn python library. Hyper parameterization of the model yielded the following good parameters: *‘learning_rate’ = 0.1, ‘max_depth’ = 1, ‘max_iter’, ‘max_leaf_nodes’ = 40, ‘min_samples_leaf’ = 20*. Hyper parameterization was conducted with 5 fold cross validation.

#### 2.3.4 Hierarchical and K-means clustering

Clustering methods cluster data together using distance values allowing us to categorize data by common attributes. Agglomerative heriarchical clusters were built from the bottom up to each of the clusters using an n of 700. K-means clusters were determined from a starting 700 centroids. Clustering was performed using gower distances from motif and node features through the stats package in R [41]. Cluster size was determined by hyperparamaterization for highest accuracy in all feature sets. Accuracy was determined by psuedolabels given based on common IDs within clusters.

#### 2.3.5 CD-Hit

CD-HIT is an incremental greedy clustering method that sorts input sequences from long to short and processes them sequentially where the first sequence is identified automatically as the first cluster representative sequence [30]. The remaining query sequences are then compared heuristically by short (3-mer to 10-mer) word search with the first sequence and grouped by ascending levels of homology into clusters. We clustered genes by thresholds ranging from 40% to 50% and assessed for the greatest accuracy. All runs were performed using 8 cores and 32 GB of memory.

#### 2.3.6 Deep Learning

We designed 8 fully connected, 7-layer networks with SoftMax activation to train and perform classification on each data set. Labels were encoded into one hot vectors and an ADAM optimizer was used to train the network by elementwise binary cross entropy loss. Over fitting was corrected for by adding Dropout and Batch normalization layers as well as weight decay through ADAM. All tests were run with hyper paramaterization using 5 fold cross validation, splitting training data sets in a 7:2:1 ratio, on 4 NVIDIA A100-SXM-80GB GPUs. Final parameters were: *’epochs’ = 25, ’batch_size’ = 256’, ’learning_rate’ =* 10^−3^, *’weight_decay’ =* 10^−5^.

### 2.4 Network Analysis

Networks were constructed from annotations in each model using the edge node lists of *E. coli* gathered with the Kegglinc R package [57]. Comparative metrics were determined using R and networks were visualized with the igraph pathway [7]. In order to calculate the topological similarity of the predicted networks to the E. coli network as a baseline, we used normalized adjacency similarity.

This similarity metric is a type of Known node-correspondence (KNC) methods ([49]) which measures the sum of equal non-zero entries in the adjacency matrices, and tries to encapsulate how structurally close to networks are. Having the same vertex ordering, this metric mainly indicates how much the of gene interactions is being predicted by our models. We used the implementation available in graph-tool package ([39].) which calculates the following value:

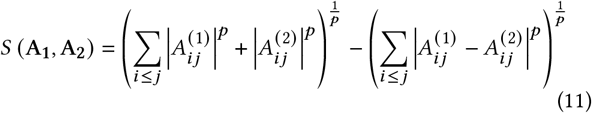

### 2.5 Data Availability

Data was collected from the KEGG database v102.0. Python and R code used in this study is available at https://github.com/RobbenUTA/ Functional-ML. Training and testing data sets have also been made available through the github repository, along with additional results for the deep learning model.

## 3 RESULTS

### 3.1 Experimental design

We developed a methodology to benchmark the performance and accuracy of machine learning models to make multi-class predictions of KEGG pathway annotations and improve the quality of pathway networks (Fig 1b). We chose to evaluate the ability for 3 ensemble, 2 clustering, and 1 deep learning model to assign function to previously annotated genes. To avoid re-annotation bias in comparison to traditional methodologies, we chose an established method of clustering-based, database alignment annotation, CD-hit, to represent homologous alignment methodologies. All machine learning models utilized the same data, with KEGG annotated genes from the genomes of *E. coli* (1,321 genes) and *B. intestinalis* (986 genes) being used for both training and validation, and genes from the genome *S. macedonicus* (583 genes) being used as a test data set.

We also compared the relative performance of homologous alignment (node) and structural sequence (motif) based feature sets. This comparison will tell us if motif or homologous sequence alignment results in better annotations. The KEGG database classifies genes by two identifiers, EC ID, biochemical and Map ID, and we determined accuracy in predicting both for each model. The EC ID groups genes by common biochemical reaction while the Map ID is typically a string of multiple identifiers that each point to a unique protein pathway in the KEGG database. We only evaluated accuracy as direct matches between predicted and real EC IDs but for Map IDs we also calculated accuracy when predicting a fraction of the Map ID numbers from the string. This allowed us to predict annotations for proteins that retain partial function when compared to a homologous gene, commonly found in truncated proteins [10].

### 3.2 Comparison of machine learning algorithms for the prediction of KEGG identifiers in training data

We determined the accuracy of each model when predicting KEGG annotations from genes in the training data set. Training data sets consisted of 1,132 genes with binary or quantitative values for 97 motifs or 3,160 nodes, and was used to determine accuracy of EC id and Map id predictions of 385 genes in a validation set. A neural network of node feature sets yielded the highest accuracy in all three class predictions (Fig. 2). Validation on the trained feature data sets suggest that homology based features are able to predict labels with greater accuracy than motif based, except for in the prediction of partial Map IDs from strings. Ensemble, clustering, and deep learning methods were shown to have increased prediction accuracy over CD-hit alignments, with Hierarchical and K Means clustering able to cluster more test genes (66% and 76% respectively) than CD-hit (10%).

**Figure 2:**
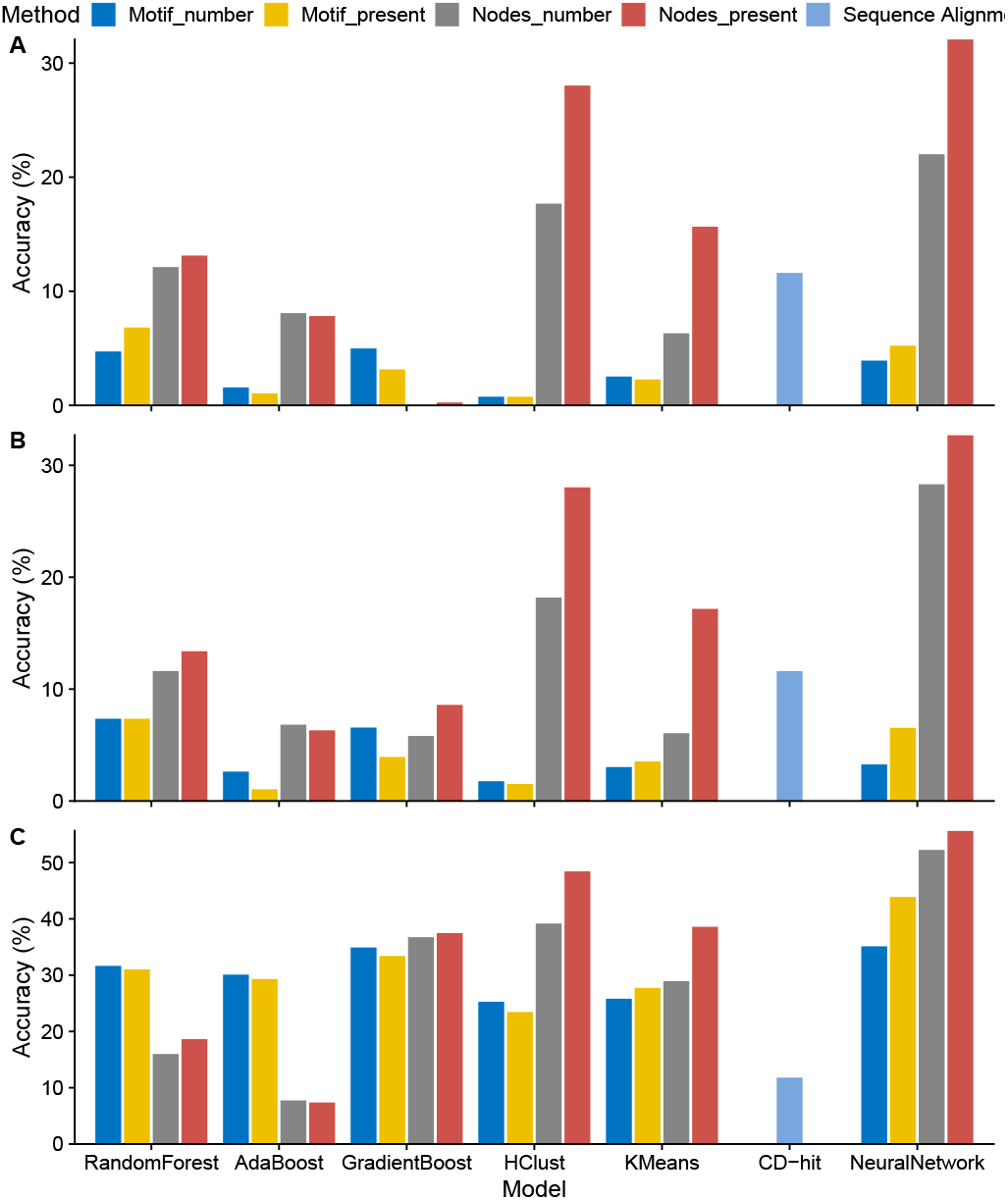
Prediction accuracy of classifier models when predicting class labels of the training data set. (A) EC ID, (B) Map ID, or (C) Partial Map feature set prediction accuracies are reported separately. Feature sets within each model were run under the same settings as determined by hyper parameterization. CD-Hit only accepts sequence information and thus was not compared for each feature set.

### 3.3 Analysis of performance metrics among models

We demonstrate that most machine learning algorithms represent an increase in speed over traditional sequence homology based methods such as CD-hit (Fig. 3a). The exception being Ada boost algorithms that took nearly 100x as long as other ensemble algorithms and 10x as long as the neural network.

**Figure 3:**
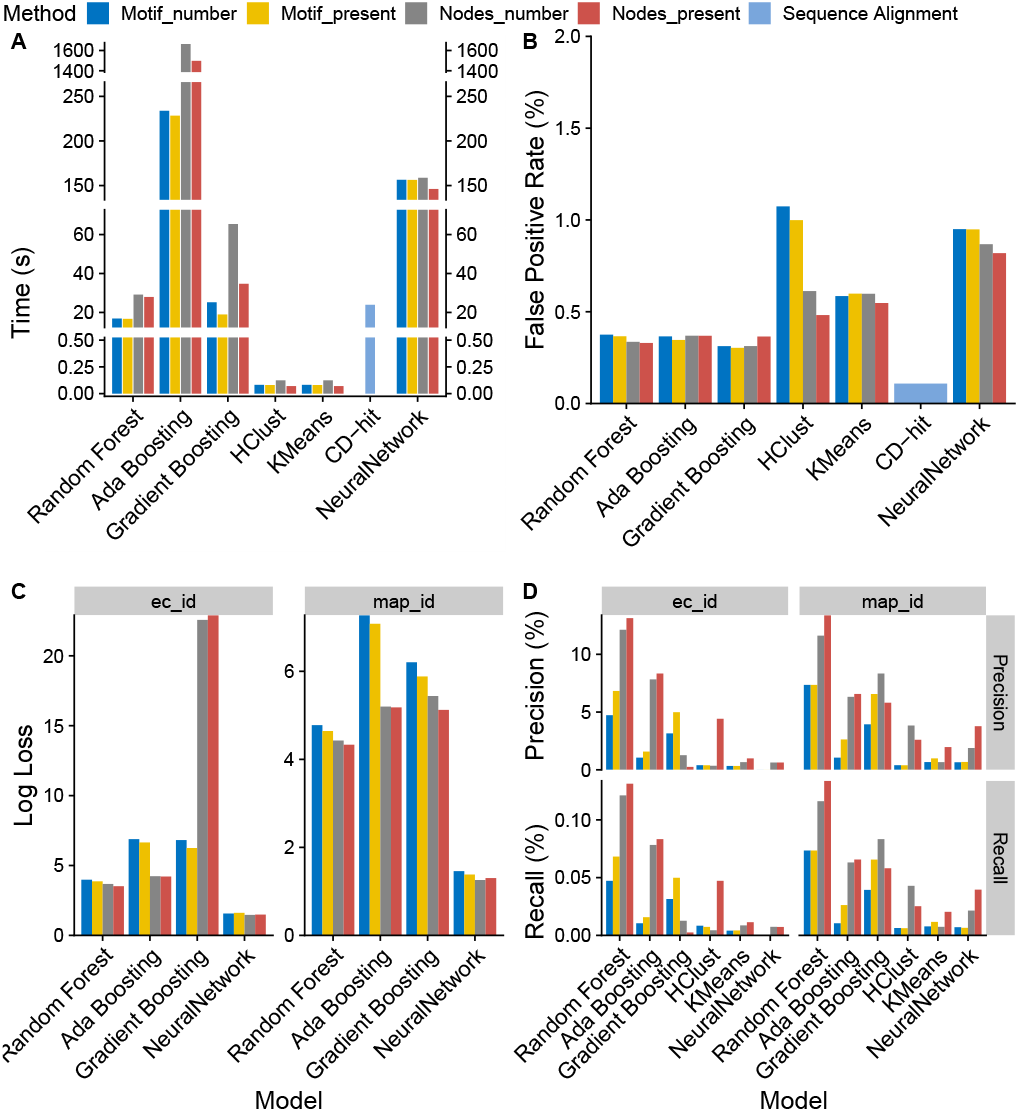
Performance and model error. (A) Computational time to train and run each machine learning model. Ensemble and clustering algorithms were all run single core, CD-Hit was multi-threaded with 8 cores and the Neural network was performed using 4 A100 GPU’s. (B) Average false positive rate among all classes reported for each model. (C) Average loss among each model, only ensemble methods and deep learning were able to report probability for each class prediction and thus loss in unsupervised methods was not shown. (D) Average precision and recall for each model was reported by percent of total gene predictions. Average precision and recall are shown for a max threshold of 100% probability only. Precision and recall was not reported for CD-hit because of low prediction resolution.

To evaluate the performance of each model, we examined various metrics of predictive power. The average False Positive Rate (FPR) among ensemble methods was fairly stagnant among feature sets and was lower than clustering and deep learning methods in every case (Fig. 3b). The surprisingly high FPR in hierarchical clustering and neural networks was not consistent with the high accuracy in the model and feature set combinations and may represent accurate predictions over fit to the training data set. Low false positive rates are reported are reported in CD-Hit, however, the algorithm only made predictions on 10% of genes.

Loss, as estimated only in ensemble and deep learning methods, was found lowest in neural networks yet is quite high across all models, potentially because only 2 species are represented in the training data set (Fig 3c). The random forest model had the highest average precision and recall across classes with both predictors (Fig 3d).

### 3.4 Motif aware machine learning models generalize predictions to new species

To validate the performance of our models on generalized data, we predicted labels from an untrained species, *S. macedonicus*. All genes in the training data set were used to predict EC id and Map id in the 583 test genes. We found that the ability for most models to generalize to a new species was fairly low, with the majority being reduced in accuracy by a factor of 10x (Fig 4). CD-hit retained high prediction accuracy in EC ID and Map ID predictions yet this did not result in increased partial Map ID prediction, presumably because it was predicting exact labels on a small subset of genes but failed to make any predictions for non-clustered genes. Every other model excluding gradient boosting yielded high partial accuracy prediction of partial Map IDs with motif data but only random forest, adaboost, and deep learning algorithms had comparable prediction accuracy with node information. Blast alignment of the test genes to those from the training data resulted in roughly 75% accuracy in all measurements, an expected result of re-annotation methods but not comparable to the machine learning algorithms used in this study, which report accuracy regardless of misannotation in the initial data set.

**Figure 4:**
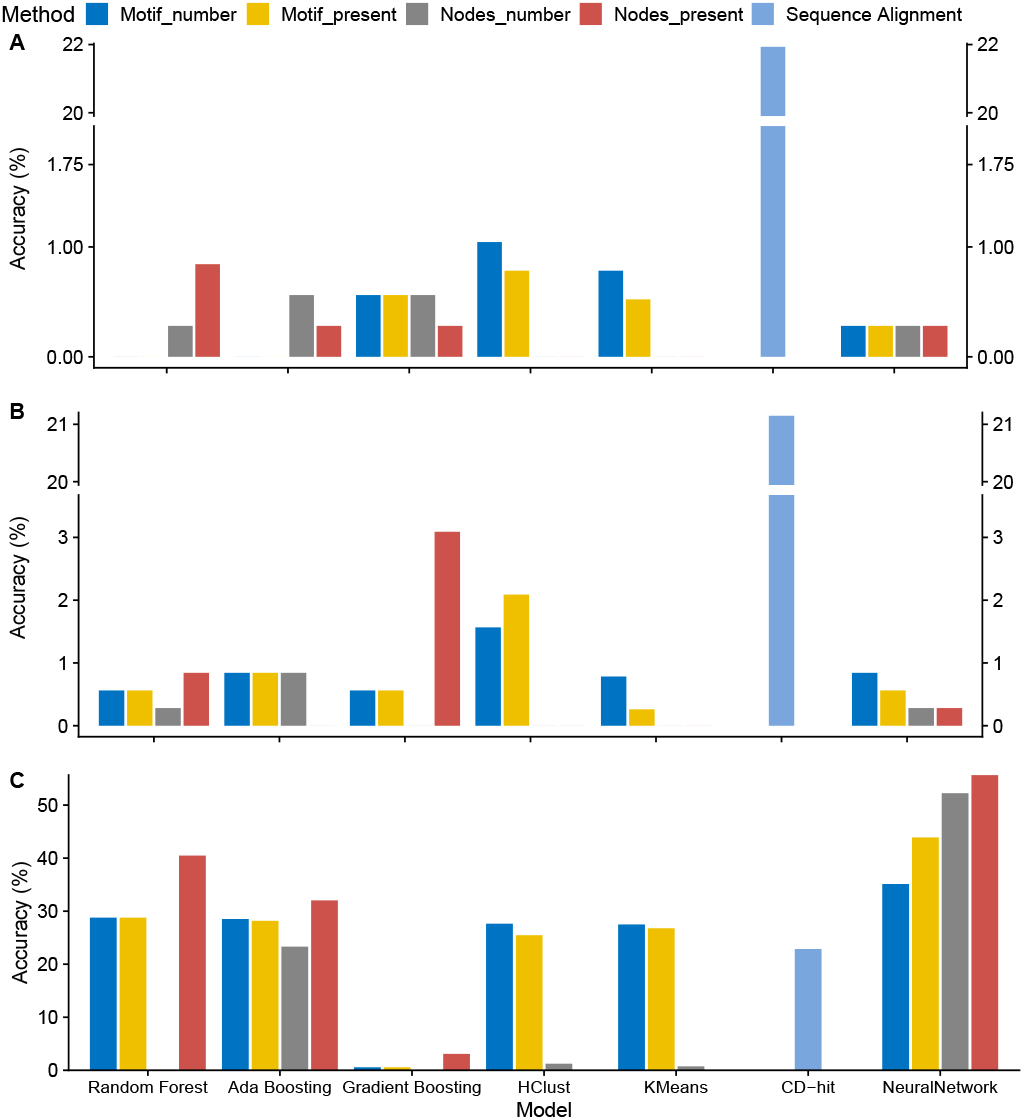
Accuracy of models to predict labels of *S. macedonicus* genes using genes from *E. coli* and *B. intestinalis* as training data. (A) EC ID, (B) Map ID, or (C) Partial Map ID feature set prediction accuracies are reported separately. Feature sets within each model were run under the same settings as determined in the previous experiment. CD-Hit only accepts sequence information and thus was not compared for each feature set. Clustering methodologies could not be trained by *E. coli* and *B. intestinalis* data separately by virtue of method.

### 3.5 Gradient Boost and Neural Network architectures improve network reconstruction of KEGG pathways

KEGG pathways are complex undirected acyclic networks constructed from biochemical evidence. As such, we can evaluate the networks reconstructed from each model annotation, and determine if the reconstructed networks lead to higher quality topology or the discovery of new pathways (Fig 5a). We found that gradient boost and neural network algorithms lead to more interconnected networks than CD-hit (Fig 5b). This was confirmed by the number of edges and level of pathway completion for each model (Fig 5c-d). These models also displayed a higher degree of similarity to *E. coli* pathways than CD-hit annotations (Fig 5e). No model had quality metrics higher than the original annotation or a BLAST alignment annotation due to a larger database with which these were annotated from. However, gradient boost and neural networks predicted more novel pathways connections than CD-hit or BLAST (Fig 5f).

**Figure 5:**
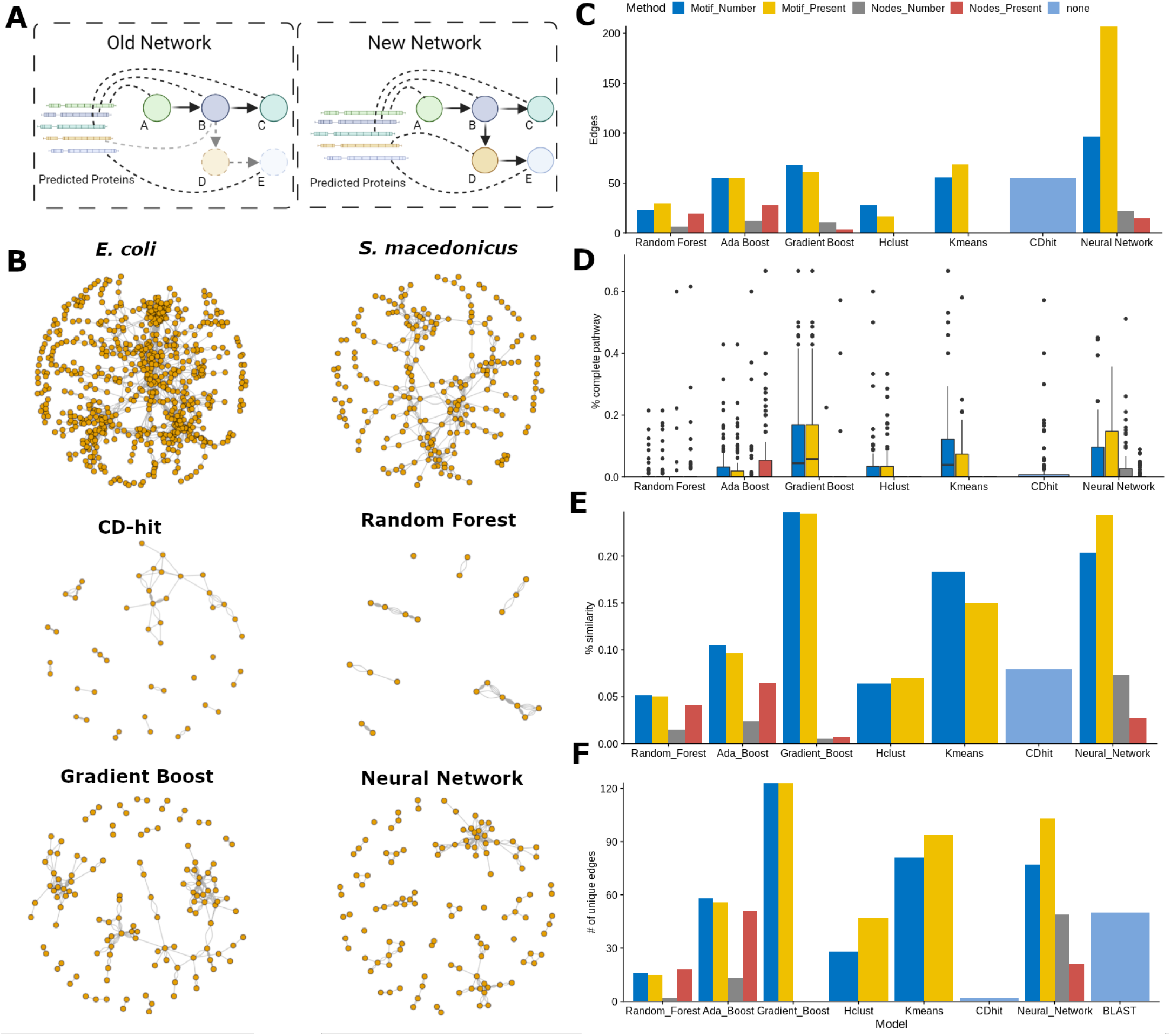
Analysis of reconstructed networks in machine learning annotations. (A) Annotation can have drastic effects on network topology, leading to the development of new metabolic and signaling pathways. Predicted protein sequences are represented as annotating to pathway nodes (circles) by dotted lines. (B) Network topology from 2 species, *E. coli* and *S. macedonicus*, and the reconstructed topologies from 4 annotation algorithms. (C) The sum of total numbers of edges and (D) the distribution of percent completion of each pathway among bench marked annotation algorithms. (E) Similarity measurements of reconstructed pathways to original pathways for each machine learning algorithm. (F) The number of unique pathway connections that were found in each machine learning model and BLAST alignment of test genes.

## 4 DISCUSSION

In this study, we sought to determine if machine learning algorithms could predict the functional annotations of bacterial genes from large scale bacterial pathway databases with higher success rates than traditional methods of sequence homology based annotation. This is an important challenge for large scale sequencing studies of the gut metagenome, as differences in metabolic pathways of gut flora have heavy ramifications on human health and disease [8]. Currently, no other study has sought to evaluate machine learning models on their ability to annotate database gene collections. Yet, related research shows an increased ability of deep learning models and SVMs to classify metagenomic read data at a higher accuracy than homologous alignment based techniques[31, 55]. Reannotation experiments using modern algorithms has proven that clustering based algorithms can solve many misannotations within databases [52].

For our experiment, we selected 7 machine learning models and one commonly used annotation methodology. Three of the models were classical ensemble machine learning algorithms which classify a test feature based on aggregate voting of labels through many individual learners. One model was a deep learning classifier that used a weighted neural network to predict labels using the same feature datasets as the ensemble methods. Two models used unsupervised learning methods to cluster the datasets by similarity to be further analyzed through pseudo-label prediction. These replicated the methods of our traditional classifier, CD-hit, which clusters orthologous genes based on sequence homology [30]. Feature dataset construction from sequence data is difficult and hampered by non-important alignment of sequence regions. We chose to utilize motifs, short regions of similar protein sequence that occur in proteins of the same function, as feature sets, either through their presence in annotated genes or combined with homology information in orthologous sequences.

While overall prediction in both validation and test data sets was rather low (<25%) for both labels, prediction of individual Map IDs from strings was far more accurate. This is likely reflective of poor gene annotation in the training data [33]. Low accuracy will also likely improve with the addition of more training data, a common effect of database size on learning models [55].

Time is an important factor for database scale annotation programs, as some commonly used sequence alignment algorithms can take weeks to process large numbers of genes. Most ML models that we tested ran as fast or faster single core than CD-hit did on multi-core tasks, and will likely also scale better (complexity (*O*)*m*) than CD-hit (complexity (*O*)*n*) because the *m* number of motifs will not increase linearly [30].

Perfect annotation of newly sequenced bacterial species is the goal of annotation algorithms but annotation quality decreases linearly with genetic distance from model species [33]. Similarly, when we validated our models on their ability to predict labels in *S. macedonicus* using *E. coli* and *B. intestinalis* genes as training data, we observed a drastic decrease in the accuracy of EC ID and Map ID predictions and reduced ability to generalize to new data. This decrease was less noticeable in partial accuracy predictions, where motif based feature sets retained consistently high results in comparison to homology based feature sets. We believe that this indicates that motif data will allow models to better generalize to new species and predict new gene function. While CD-hit had consistent accuracy across all predictors, the model only predicted function in 25% of genes as compared to 100% in ensemble and deep learning models and about 80% in clustering methods. Previous implementations of CD-hit also observed a concerningly high lack of annotation in new gene sets [52].

The most interesting aspect of this study is how each machine learning algorithm effects the quality of the KEGG network. As we demonstrate in Fig 5a, reconstruction of biological networks can lead to the discovery of new biology. Correctly annotating predicted proteins can determine the kinds of metabolites a species can process or connect entire parts of pathway networks together. In our case, we were able to find new pathways that did not exist in the original annotation when classifying genes by gradient boosting and neural network architectures. This improved connectivity correlated with general measures of pathway connectivity and increased node degrees. Because of their application to non-curated data, reconstructed networks can be used for the elucidation of new biology, such as the discovery of new species and metabolic pathways in gut microbiome samples [60]. Reconstructed networks can also be created from individually sequenced faecal samples to catch incidents of horizontal gene transfer and random mutation that lead to novel metabolic function [51].

Due of its ability to retain high accuracy and network connectivity when used to predict generalized classifications of gene function, neural networks represent the best use case of machine learning for this application. Other models had admirable aspects as well, with random forests producing the lowest FPR and highest precision-recall, and gradient boosting algorithms classifying the greatest amount of novel pathway function. The choice from this benchmark as to which algorithm to apply to a new tool would depend highly on the intended purpose of the tool. Whether the priority is recreation of network topology and high accuracy or the discovery of new biology. We believe that the results presented in this study will assist future researchers in determining the best algorithm to use with their tools.

The machine learning algorithms we investigated in this study and the networks that can be reconstructed from them can be applied to many practical uses in network computational biology. Obviously, database quality is a major concern for many researchers as it affects the quality of experiments that rely on good annotations [40, 47]. Many integrated pipelines to annotate newly sequenced genomes restrict the researchers ability to use new or better annotation methods to annotate genomes prior to submission to databases [21, 35, 50]. Also, annotation of non-model bacterial genomes is highly sensitive to the data in which it is annotated from, and higher accuracy in predictions is a priority for these experiments, especially in the case of horizontal gene transfer between gut bacteria [11, 32, 43, 48]. Genetic screening of human metabolic pathways in disease progression is one field which could benefit from better annotation of metabolic genes [61]. Using machine learning assisted annotation we can develop pipelines to investigate bacterial pathway effects on human health, which will increase the amount of data that can be applied to large scale networks of pathways and diseases (Fig 6). Metabolic pathway screening for gut microbiomes requires high fidelity gene annotations [42] and this problem cannot be solved without annotation methodologies that more accurately recreate function. We have shown that motif based machine learning classification of gene functional annotation in two human gut microbes, *B. intestinalis* and *S. macedonicus*, results in higher accuracy and better network reconstruction when compared to functional annotation by equivalent sequence alignment technology. From what we have shown in this study, machine learning algorithms can be used to improve networks that rely on annotation of sequenced genomes. Our results represent a powerful next step in utilizing machine learning to solve the problems of network construction from sequencing data in biological databases.

**Figure 6:**
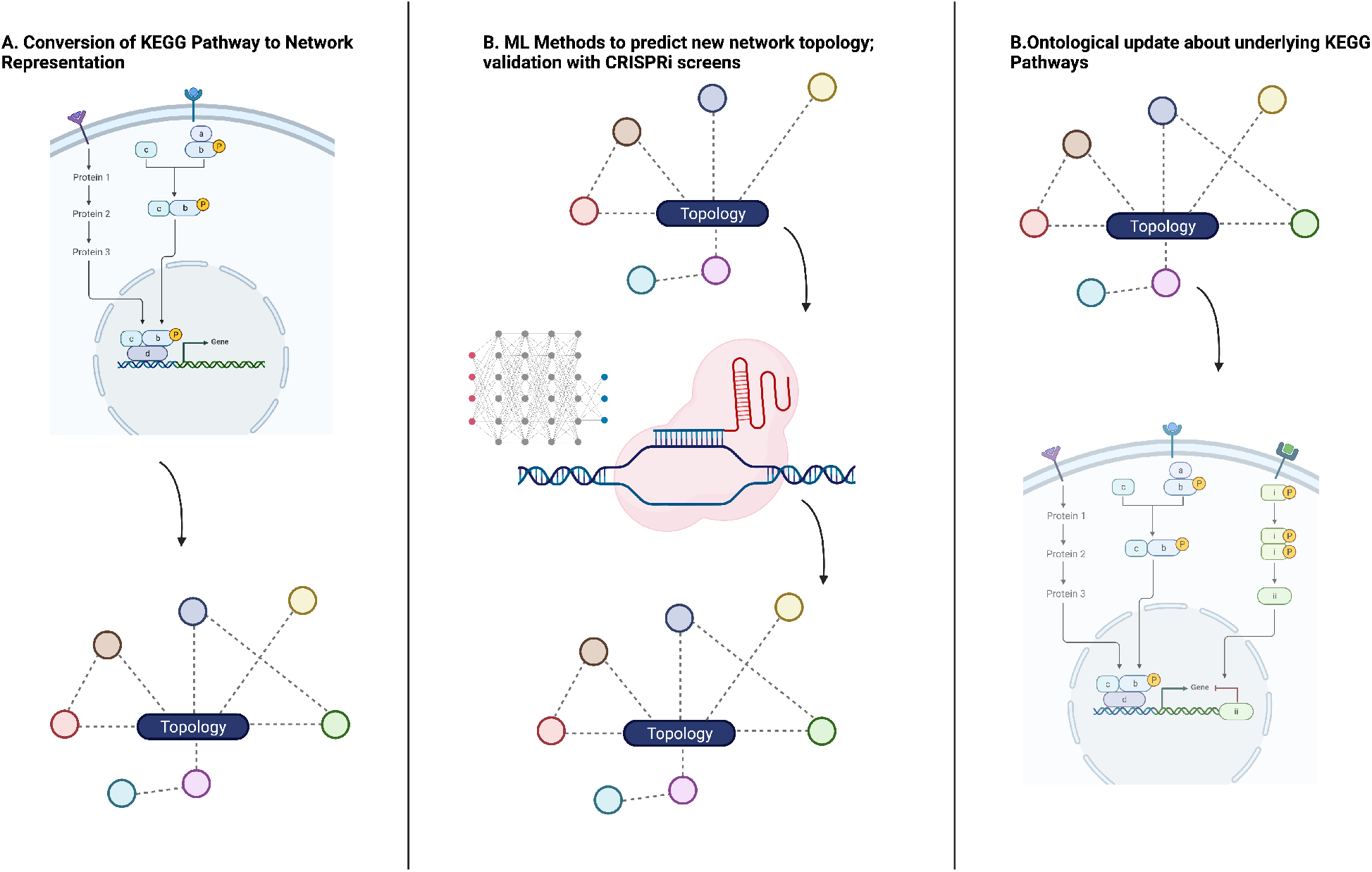
Pipeline for the discovery of bacterial metabolic pathways in the effects on human health.

## ACKNOWLEDGMENTS

The published work was made possible by the University of Texas System Rising STARs Award (J.M.L) and the CPRIT First Time Faculty Award (J.M.L). The authors would also like to express their thanks to Fiza Saeed for her assistance with the curation of data from the KEGG database.

## REFERENCES

[1] Xiangning Bai, Aswathy Narayanan, Piotr Nowak, Shilpa Ray, Ujjwal Neogi, and Anders Sönnerborg. 2021. Whole-Genome Metagenomic Analysis of the Gut Microbiome in HIV-1-Infected Individuals on Antiretroviral Therapy. Front. Microbiol. 12 (June 2021), 667718.

[2] Timothy L Bailey, James Johnson, Charles E Grant, and William S Noble. 2015. The MEME Suite. Nucleic Acids Res. 43, W1 (July 2015), W39–49.

[3] Jacob Beal, Adam Clore, and Jeff Manthey. 2022. Studying Pathogens Degrades BLAST-based Pathogen Identification. bioRxiv (2022).

[4] Matthias Blum, Hsin-Yu Chang, Sara Chuguransky, Tiago Grego, Swaathi Kandasaamy, Alex Mitchell, Gift Nuka, Typhaine Paysan-Lafosse, Matloob Qureshi, Shriya Raj, Lorna Richardson, Gustavo A Salazar, Lowri Williams, Peer Bork, Alan Bridge, Julian Gough, Daniel H Haft, Ivica Letunic, Aron Marchler-Bauer, Huaiyu Mi, Darren A Natale, Marco Necci, Christine A Orengo, Arun P Pandurangan, Catherine Rivoire, Christian J A Sigrist, Ian Sillitoe, Narmada Thanki, Paul D Thomas, Silvio C E Tosatto, Cathy H Wu, Alex Bateman, and Robert D Finn. 2021. The InterPro protein families and domains database: 20 years on. Nucleic Acids Res. 49, D1 (Jan. 2021), D344-D354.

[5] Leo Breiman. 2001. Random Forests. Mach. Learn. 45, 1 (Oct. 2001), 5–32.

[6] Benjamin Buchfink, Chao Xie, and Daniel H Huson. 2015. Fast and sensitive protein alignment using DIAMOND. Nat. Methods 12, 1 (Jan. 2015), 59–60.

[7] Gabor Csardi and Tamas Nepusz. 2006. The igraph software package for complex network research. InterJournal Complex Systems (2006), 1695. https://igraph.org

[8] Yong Fan and Oluf Pedersen. 2021. Gut microbiota in human metabolic health and disease. Nat. Rev. Microbiol. 19, 1 (Jan. 2021), 55–71.

[9] Robert D Finn, Jody Clements, and Sean R Eddy. 2011. HMMER web server: interactive sequence similarity searching. Nucleic Acids Res. 39, Web Server issue (July 2011), W29–37.

[10] Nikolaus Fortelny, Paul Pavlidis, and Christopher M Overall. 2015. The path of no return—Truncated protein N-termini and current ignorance of their genesis. Proteomics 15, 14 (2015), 2547–2552.

[11] Eric A Franzosa, Lauren J McIver, Gholamali Rahnavard, Luke R Thompson, Melanie Schirmer, George Weingart, Karen Schwarzberg Lipson, Rob Knight, J Gregory Caporaso, Nicola Segata, and Curtis Huttenhower. 2018. Species-level functional profiling of metagenomes and metatranscriptomes. Nat. Methods 15, 11 (Nov. 2018), 962–968.

[12] Yoav Freund and Robert E Schapire. 1997. A Decision-Theoretic Generalization of On-Line Learning and an Application to Boosting. J. Comput. System Sci. 55, 1 (Aug. 1997), 119–139.

[13] Jerome H Friedman. 2002. Stochastic gradient boosting. Comput. Stat. Data Anal. 38, 4 (Feb. 2002), 367–378.

[14] María José Gosalbes, Ana Durbán, Miguel Pignatelli, Juan José Abellan, Nuria Jiménez-Hernández, Ana Elena Pérez-Cobas, Amparo Latorre, and Andrés Moya. 2011. Metatranscriptomic approach to analyze the functional human gut microbiota. PLoS One 6, 3 (March 2011), e17447.

[15] Charles E Grant, Timothy L Bailey, and William Stafford Noble. 2011. FIMO: scanning for occurrences of a given motif. Bioinformatics 27, 7 (April 2011), 1017–1018.

[16] M L Green and P D Karp. 2005. Genome annotation errors in pathway databases due to semantic ambiguity in partial EC numbers. Nucleic Acids Res. 33, 13 (July 2005), 4035–4039.

[17] Margaret G Guo, Daniel N Sosa, and Russ B Altman. 2022. Challenges and opportunities in network-based solutions for biological questions. Briefings in Bioinformatics 23, 1 (2022), bbab437.

[18] Beth A Helmink, M A Wadud Khan, Amanda Hermann, Vancheswaran Gopalakrishnan, and Jennifer A Wargo. 2019. The microbiome, cancer, and cancer therapy. Nat. Med. 25, 3 (March 2019), 377–388.

[19] Human Microbiome Project Consortium. 2012. Structure, function and diversity of the healthy human microbiome. Nature 486, 7402 (June 2012), 207–214.

[20] Sushrut Jangi, Roopali Gandhi, Laura M Cox, Ning Li, Felipe von Glehn, Raymond Yan, Bonny Patel, Maria Antonietta Mazzola, Shirong Liu, Bonnie L Glanz, Sandra Cook, Stephanie Tankou, Fiona Stuart, Kirsy Melo, Parham Nejad, Kathleen Smith, Begüm D Topçuolu, James Holden, Pia Kivisäkk, Tanuja Chitnis, Philip L De Jager, Francisco J Quintana, Georg K Gerber, Lynn Bry, and Howard L Weiner. 2016. Alterations of the human gut microbiome in multiple sclerosis. Nat. Commun. 7 (June 2016), 12015.

[21] Philip Jones, David Binns, Hsin-Yu Chang, Matthew Fraser, Weizhong Li, Craig McAnulla, Hamish McWilliam, John Maslen, Alex Mitchell, Gift Nuka, et al. 2014. InterProScan 5: genome-scale protein function classification. Bioinformatics 30, 9 (2014), 1236–1240.

[22] M Kanehisa and S Goto. 2000. KEGG: kyoto encyclopedia of genes and genomes. Nucleic Acids Res. 28, 1 (Jan. 2000), 27–30.

[23] Kazutaka Katoh, Kazuharu Misawa, Kei-Ichi Kuma, and Takashi Miyata. 2002. MAFFT: a novel method for rapid multiple sequence alignment based on fast Fourier transform. Nucleic Acids Res. 30, 14 (July 2002), 3059–3066.

[24] Mohammed Khalilia, Sounak Chakraborty, and Mihail Popescu. 2011. Predicting disease risks from highly imbalanced data using random forest. BMC Med. Inform. Decis. Mak. 11 (July 2011), 51.

[25] Donghyun Kim, Melody Y Zeng, and Gabriel Núñez. 2017. The interplay between host immune cells and gut microbiota in chronic inflammatory diseases. Exp. Mol. Med. 49, 5 (May 2017), e339.

[26] Toshihiro Kishikawa, Yuichi Maeda, Takuro Nii, Daisuke Motooka, Yuki Matsumoto, Masato Matsushita, Hidetoshi Matsuoka, Maiko Yoshimura, Shoji Kawada, Satoru Teshigawara, Eri Oguro, Yasutaka Okita, Keisuke Kawamoto, Shinji Higa, Toru Hirano, Masashi Narazaki, Atsushi Ogata, Yukihiko Saeki, Shota Nakamura, Hidenori Inohara, Atsushi Kumanogoh, Kiyoshi Takeda, and Yukinori Okada. 2020. Metagenome-wide association study of gut microbiome revealed novel aetiology of rheumatoid arthritis in the Japanese population. Ann. Rheum. Dis. 79, 1 (Jan. 2020), 103–111.

[27] Toshihiro Kishikawa, Kotaro Ogawa, Daisuke Motooka, Akiko Hosokawa, Makoto Kinoshita, Ken Suzuki, Kenichi Yamamoto, Tatsuo Masuda, Yuki Matsumoto, Takuro Nii, Yuichi Maeda, Shota Nakamura, Hidenori Inohara, Hideki Mochizuki, Tatsusada Okuno, and Yukinori Okada. 2020. A Metagenome-Wide Association Study of Gut Microbiome in Patients With Multiple Sclerosis Revealed Novel Disease Pathology. Front. Cell. Infect. Microbiol. 10 (Dec. 2020), 585973.

[28] Weizhong Li. 2009. Analysis and comparison of very large metagenomes with fast clustering and functional annotation. BMC Bioinformatics 10 (Oct. 2009), 359.

[29] Weizhong Li, Limin Fu, Beifang Niu, Sitao Wu, and John Wooley. 2012. Ultrafast clustering algorithms for metagenomic sequence analysis. Brief. Bioinform. 13, 6 (Nov. 2012), 656–668.

[30] Weizhong Li and Adam Godzik. 2006. Cd-hit: a fast program for clustering and comparing large sets of protein or nucleotide sequences. Bioinformatics 22, 13 (July 2006), 1658–1659.

[31] Qiaoxing Liang, Paul W Bible, Yu Liu, Bin Zou, and Lai Wei. 2020. DeepMicrobes: taxonomic classification for metagenomics with deep learning. NAR Genom Bioinform 2, 1 (March 2020), qaa009.

[32] Jason Lloyd-Price, Anup Mahurkar, Gholamali Rahnavard, Jonathan Crabtree, Joshua Orvis, A Brantley Hall, Arthur Brady, Heather H Creasy, Carrie McCracken, Michelle G Giglio, Daniel McDonald, Eric A Franzosa, Rob Knight, Owen White, and Curtis Huttenhower. 2017. Strains, functions and dynamics in the expanded Human Microbiome Project. Nature 550, 7674 (Oct. 2017), 61–66.

[33] Briallen Lobb, Benjamin Jean-Marie Tremblay, Gabriel Moreno-Hagelsieb, and Andrew C Doxey. 2020. An assessment of genome annotation coverage across the bacterial tree of life. Microb Genom 6, 3 (March 2020).

[34] Jaina Mistry, Sara Chuguransky, Lowri Williams, Matloob Qureshi, Gustavo A Salazar, Erik L L Sonnhammer, Silvio C E Tosatto, Lisanna Paladin, Shriya Raj, Lorna J Richardson, Robert D Finn, and Alex Bateman. 2021. Pfam: The protein families database in 2021. Nucleic Acids Res. 49, D1 (Jan. 2021), D412–D419.

[35] Yuki Moriya, Masumi Itoh, Shujiro Okuda, Akiyasu C Yoshizawa, and Minoru Kanehisa. 2007. KAAS: an automatic genome annotation and pathway reconstruction server. Nucleic acids research 35, suppl_2 (2007), W182–W185.

[36] Tania Nobre, M Doroteia Campos, Eva Lucic-Mercy, and Birgit Arnholdt-Schmitt. 2016. Misannotation Awareness: A Tale of Two Gene-Groups. Front. Plant Sci. 7 (June 2016), 868.

[37] Nuala A O’Leary, Mathew W Wright, J Rodney Brister, Stacy Ciufo, Diana Haddad, Rich McVeigh, Bhanu Rajput, Barbara Robbertse, Brian Smith-White, Danso Ako-Adjei, Alexander Astashyn, Azat Badretdin, Yiming Bao, Olga Blinkova, Vyacheslav Brover, Vyacheslav Chetvernin, Jinna Choi, Eric Cox, Olga Ermolaeva, Catherine M Farrell, Tamara Goldfarb, Tripti Gupta, Daniel Haft, Eneida Hatcher, Wratko Hlavina, Vinita S Joardar, Vamsi K Kodali, Wenjun Li, Donna Maglott, Patrick Masterson, Kelly M McGarvey, Michael R Murphy, Kathleen O’Neill, Shashikant Pujar, Sanjida H Rangwala, Daniel Rausch, Lillian D Riddick, Conrad Schoch, Andrei Shkeda, Susan S Storz, Hanzhen Sun, Francoise Thibaud-Nissen, Igor Tolstoy, Raymond E Tully, Anjana R Vatsan, Craig Wallin, David Webb, Wendy Wu, Melissa J Landrum, Avi Kimchi, Tatiana Tatusova, Michael DiCuccio, Paul Kitts, Terence D Murphy, and Kim D Pruitt. 2016. Reference sequence (RefSeq) database at NCBI: current status, taxonomic expansion, and functional annotation. Nucleic Acids Res. 44, D1 (Jan. 2016), D733–45.

[38] Ross Overbeek, Tadhg Begley, Ralph M Butler, Jomuna V Choudhuri, Han-Yu Chuang, Matthew Cohoon, Valérie de Crécy-Lagard, Naryttza Diaz, Terry Disz, Robert Edwards, Michael Fonstein, Ed D Frank, Svetlana Gerdes, Elizabeth M Glass, Alexander Goesmann, Andrew Hanson, Dirk Iwata-Reuyl, Roy Jensen, Neema Jamshidi, Lutz Krause, Michael Kubal, Niels Larsen, Burkhard Linke, Alice C McHardy, Folker Meyer, Heiko Neuweger, Gary Olsen, Robert Olson, Andrei Osterman, Vasiliy Portnoy, Gordon D Pusch, Dmitry A Rodionov, Christian Rückert, Jason Steiner, Rick Stevens, Ines Thiele, Olga Vassieva, Yuzhen Ye, Olga Zagnitko, and Veronika Vonstein. 2005. The subsystems approach to genome annotation and its use in the project to annotate 1000 genomes. Nucleic Acids Res. 33, 17 (Oct. 2005), 5691–5702.

[39] Tiago P. Peixoto. 2014. The graph-tool python library. figshare (2014). https://doi.org/10.6084/m9.figshare.1164194

[40] Junjie Qin, Ruiqiang Li, Jeroen Raes, Manimozhiyan Arumugam, Kristoffer Solvsten Burgdorf, Chaysavanh Manichanh, Trine Nielsen, Nicolas Pons, Florence Levenez, Takuji Yamada, Daniel R Mende, Junhua Li, Junming Xu, Shaochuan Li, Dongfang Li, Jianjun Cao, Bo Wang, Huiqing Liang, Huisong Zheng, Yinlong Xie, Julien Tap, Patricia Lepage, Marcelo Bertalan, Jean-Michel Batto, Torben Hansen, Denis Le Paslier, Allan Linneberg, H Bjørn Nielsen, Eric Pelletier, Pierre Renault, Thomas Sicheritz-Ponten, Keith Turner, Hongmei Zhu, Chang Yu, Shengting Li, Min Jian, Yan Zhou, Yingrui Li, Xiuqing Zhang, Songgang Li, Nan Qin, Huanming Yang, Jian Wang, Søren Brunak, Joel Doré, Francisco Guarner, Karsten Kristiansen, Oluf Pedersen, Julian Parkhill, Jean Weissenbach, MetaHIT Consortium, Peer Bork, S Dusko Ehrlich, and Jun Wang. 2010. A human gut microbial gene catalogue established by metagenomic sequencing. Nature 464, 7285 (March 2010), 59–65.

[41] R Core Team. 2013. R: A Language and Environment for Statistical Computing. R Foundation for Statistical Computing, Vienna, Austria. http://www.R-project.org/ ISBN 3-900051-07-0.

[42] François Rousset and David Bikard. 2020. CRISPR screens in the era of microbiomes. Curr. Opin. Microbiol. 57 (Oct. 2020), 70–77.

[43] Chicago, IL Steven L Salzberg. 2019. Next-generation genome annotation: we still struggle to get it right. Genome Biol. 20, 1 (May 2019), 92.

[44] R Sandberg, G Winberg, C I Bränden, A Kaske, I Ernberg, and J Cöster. 2001. Capturing whole-genome characteristics in short sequences using a naïve Bayesian classifier. Genome Res. 11, 8 (Aug. 2001), 1404–1409.

[45] Eric W Sayers, Evan E Bolton, J Rodney Brister, Kathi Canese, Jessica Chan, Donald C Comeau, Ryan Connor, Kathryn Funk, Chris Kelly, Sunghwan Kim, Tom Madej, Aron Marchler-Bauer, Christopher Lanczycki, Stacy Lathrop, Zhiyong Lu, Francoise Thibaud-Nissen, Terence Murphy, Lon Phan, Yuri Skripchenko, Tony Tse, Jiyao Wang, Rebecca Williams, Barton W Trawick, Kim D Pruitt, and Stephen T Sherry. 2022. Database resources of the national center for biotechnology information. Nucleic Acids Res. 50, D1 (Jan. 2022), D20–D26.

[46] Klaus Peter Schliep. 2011. phangorn: phylogenetic analysis in R. Bioinformatics 27, 4 (Feb. 2011), 592–593.

[47] Alexandra M Schnoes, Shoshana D Brown, Igor Dodevski, and Patricia C Babbitt. 2009. Annotation error in public databases: misannotation of molecular function in enzyme superfamilies. PLoS Comput. Biol. 5, 12 (Dec. 2009), e1000605.

[48] Torsten Seemann. 2014. Prokka: rapid prokaryotic genome annotation. Bioinformatics 30, 14 (July 2014), 2068–2069.

[49] Mattia Tantardini, Francesca Ieva, Lucia Tajoli, and Carlo Piccardi. 2019. Comparing methods for comparing networks. Sci. Rep. 9, 1 (Nov. 2019), 17557.

[50] Tatiana Tatusova, Michael DiCuccio, Azat Badretdin, Vyacheslav Chetvernin, Eric P Nawrocki, Leonid Zaslavsky, Alexandre Lomsadze, Kim D Pruitt, Mark Borodovsky, and James Ostell. 2016. NCBI prokaryotic genome annotation pipeline. Nucleic Acids Res. 44, 14 (Aug. 2016), 6614–6624.

[51] Braden T Tierney, Erika Szymanski, James R Henriksen, Aleksandar D Kostic, and Chirag J Patel. 2021. Using Cartesian Doubt To Build a Sequencing-Based View of Microbiology. Msystems 6, 5 (2021), e00574–21.

[52] Braden T Tierney, Zhen Yang, Jacob M Luber, Marc Beaudin, Marsha C Wibowo, Christina Baek, Eleanor Mehlenbacher, Chirag J Patel, and Aleksandar D Kostic. 2019. The Landscape of Genetic Content in the Gut and Oral Human Microbiome. Cell Host Microbe 26, 2 (Aug. 2019), 283–295.e8.

[53] Shahadat Uddin, Arif Khan, Md Ekramul Hossain, and Mohammad Ali Moni. 2019. Comparing different supervised machine learning algorithms for disease prediction. BMC Med. Inform. Decis. Mak. 19, 1 (Dec. 2019), 281.

[54] UniProt Consortium. 2021. UniProt: the universal protein knowledgebase in 2021. Nucleic Acids Res. 49, D1 (Jan. 2021), D480–D489.

[55] Kévin Vervier, Pierre Mahé, Maud Tournoud, Jean-Baptiste Veyrieras, and Jean-Philippe Vert. 2016. Large-scale machine learning for metagenomics sequence classification. Bioinformatics 32, 7 (April 2016), 1023–1032.

[56] Xi Wang, Yujie Ning, Cheng Li, Yi Gong, Ruitian Huang, Minhan Hu, Blandine Poulet, Ke Xu, Guanghui Zhao, Rong Zhou, Mikko J Lammi, and Xiong Guo. 2021. Alterations in the gut microbiota and metabolite profiles of patients with Kashin-Beck disease, an endemic osteoarthritis in China. Cell Death Dis. 12, 11 (Oct. 2021), 1015.

[57] Shana White and Mario Medvedovic. 2016. KEGGlincs design and application: an R package for exploring relationships in biological pathways [version 1; not peer reviewed]. https://doi.org/10.7490/f1000research.1113436.1

[58] Derrick E Wood, Jennifer Lu, and Ben Langmead. 2019. Improved metagenomic analysis with Kraken 2. Genome Biol. 20, 1 (Nov. 2019), 257.

[59] Guangchuang Yu. 2022. tidytree: A Tidy Tool for Phylogenetic Tree Data Manipulation. https://yulab-smu.top/treedata-book/ R package version 0.3.9.

[60] Tatyana Zamkovaya, Jamie S Foster, Valérie de Crécy-Lagard, and Ana Conesa. 2021. A network approach to elucidate and prioritize microbial dark matter in microbial communities. The ISME journal 15, 1 (2021), 228–244.

[61] Dongxin Zhao, Mehmet G Badur, Jens Luebeck, Jose H Magaña, Amanda Birmingham, Roman Sasik, Christopher S Ahn, Trey Ideker, Christian M Metallo, and Prashant Mali. 2018. Combinatorial CRISPR-Cas9 Metabolic Screens Reveal Critical Redox Control Points Dependent on the KEAP1-NRF2 Regulatory Axis. Mol. Cell 69, 4 (Feb. 2018), 699–708.e7.

[62] Laurence Zitvogel, Romain Daillère, María Paula Roberti, Bertrand Routy, and Guido Kroemer. 2017. Anticancer effects of the microbiome and its products. Nat. Rev. Microbiol. 15, 8 (Aug. 2017), 465–478.

